# Extinction models of robustness for weighted ecological networks

**DOI:** 10.1101/186577

**Authors:** Miranda S. Bane, Michael J. O. Pocock, Richard James

## Abstract

1. Analysis of ecological networks is a valuable approach to understanding the vulnerability of systems to environmental change. The tolerance of ecological networks to co-extinctions, resulting from sequences of primary extinctions, is a widely-used tool for modelling network ‘robustness’. Previously, these ‘extinction models’ have been developed for and applied mostly to binary networks and recently used to predict cascades of co-extinctions in plant-pollinator networks. There is a need for robustness models that can make the most of the weighted data available and most importantly there is a need to understand how the structure of a network affects its robustness.
2. Here, we developed a framework of extinction models for bipartite ecological networks (specifically plant-pollinator networks). In previous models co-extinctions occurred when nodes lost all their links, but by relaxing this rule (according to a set threshold) our models can be applied to binary and weighted networks, and can permit structurally correlated extinctions, i.e. the potential for avalanches of extinctions. We tested how the average and the range of robustness values is impacted by network structure and the impact of structurally-correlated extinctions sampling non-uniformly from the distribution of random extinction sequences.
3. We found that the way that structurally-correlated extinctions are modelled impacts the results; our two ecologically-plausible models produce opposing effects which shows the importance of understanding the model. We found that when applying the models to networks with weighted interactions, the effects are amplified and the variation in robustness increases. Variation in robustness is a key feature of these extinction models and is driven by the structural heterogeneity (i.e. the skewness of the degree distribution) of nodes (specifically, plant nodes) in the network.
4. Our new framework of models enables us to calculate robustness with weighted, as well as binary, bipartite networks, and to make direct comparisons between models and between networks. This allows us to differentiate effects of the model and of the data (network structure) which is vital for those making ecological inferences from robustness models. The models can be applied to mutualistic and antagonistic networks, and can be extended to food webs.

## 1. INTRODUCTION

Network analysis has become an important tool for ecologists seeking to understand the vulnerability of ecosystems to environmental change. Recent research has centred on network approaches for improving our understanding of plant-pollinator communities and extinctions, especially in the light of the widely documented recent declines in key insect pollinators such as honeybees, bumblebees and butterflies (Biesmeijer *et al.,* 2006; Senapathi *et al.*, 2015; Goulson, Lye & Darvill, 2008; Benton, 2006). These trends are concerning for biodiversity, ecosystem function and food security (Potts *et al*., 2010) as insect pollinators are known to play a vital role in providing ecosystem services (Bailes *et al.*, 2015). They feed on nectar and pollen provided by plant species, and whilst doing this facilitate the fertilisation of plants via cross pollination (Free, 1993; Lubbock, 1875). In many ecological systems, including plant-pollinator systems, the community can be regarded as a bipartite network comprising two distinct guilds of organisms in which each node represents a species, and species are connected by edges indicating interactions, which may be directly observed, indirectly observed (e.g. pollen analysis) or inferred (Morales-Castilla *et al.* 2015).

Models of community robustness based on observed plant-pollinator networks (available, for example, from http://www.web-of-life.es and https://www.nceas.ucsb.edu/interactionweb/resources.html) usually fall into one of two types. In the first (see for example Bastolla *et al.*, 2009, James, Pitchford and Plank, 2012), the community is modelled as a dynamical system, in which the population of each species is affected by the interactions that species has with others. The dynamics are typically run to fixation, and the populations at fixation used to determine community robustness, and how it relates to overall network structure.

The second approach, which we adopt here, is to model the tolerance of the network to extinctions. This was pioneered by Albert, Jeong & Barabasi (2000) and rapidly applied first to multitrophic food webs (Dunne, Williams and Martinez, 2002) and then mutualistic bipartite networks, especially plant-pollinator networks (Memmott, Waser and Price, 2004; Kaiser-Bunbury *et al.,* 2010).

Extinction models estimate the robustness of a plant-pollinator network by sequentially removing species of the primary type (e.g. plants) and recording the number of surviving species of the secondary type (e.g. pollinators), by applying some pre-determined rule for species survival. Most models, thus far, have used simple rules for secondary extinctions e.g. species become extinct when all their existing links are lost. Network robustness can then be determined from the area under the curve of the proportion of the secondary type that survive against the proportion of the primary type removed (Burgos *et al.,* 2007; see Fig. 1a).

**Figure 1:**
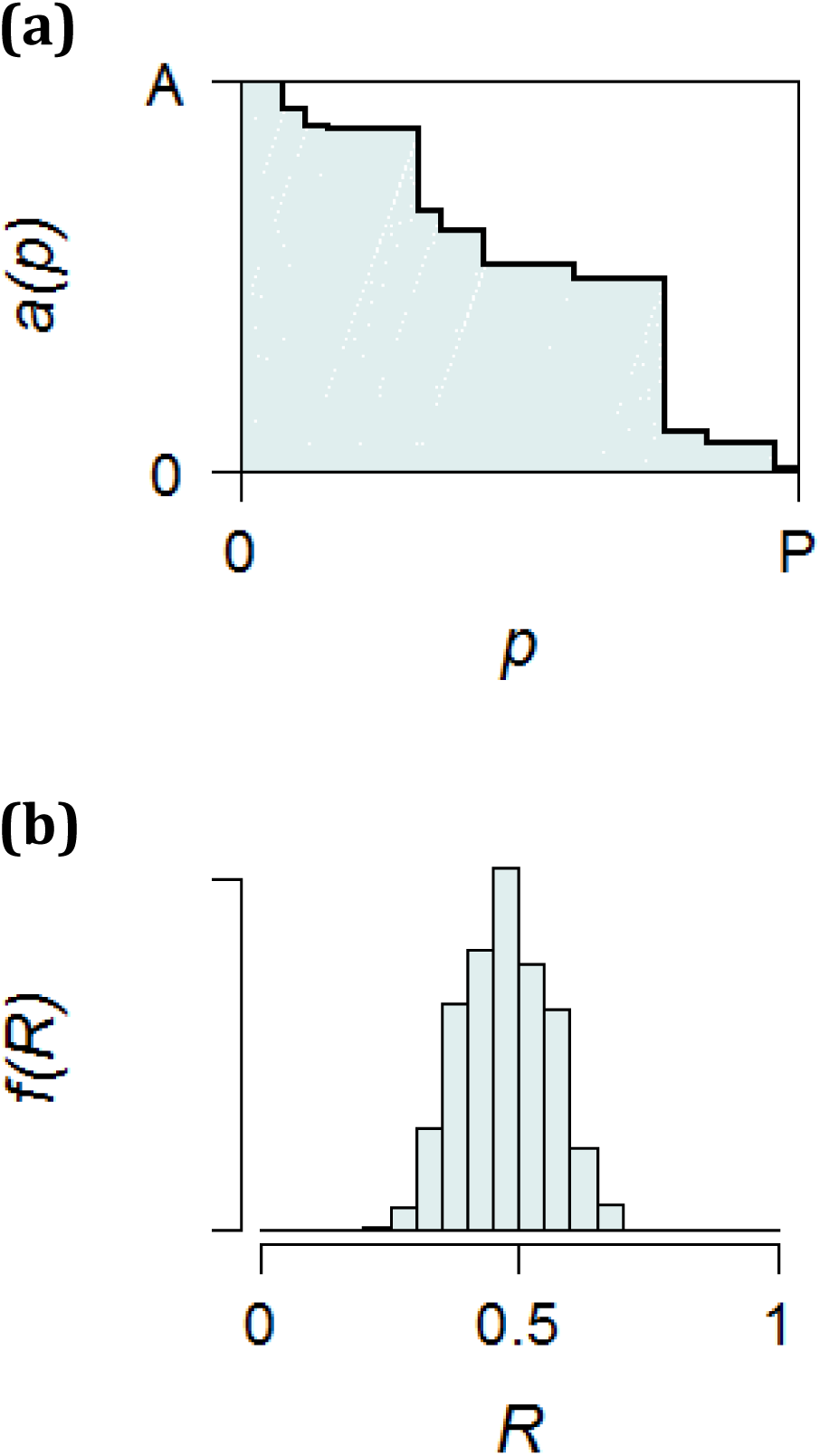
The output of an extinction model**. (a)** For a single extinction sequence the number of surviving pollinator nodes *a* reduces as the number of plant nodes made extinct, *p*, increases. There are no surviving nodes at the end of the sequence. Robustness (*R*) = 0.5504 is computed as the area under *a*(*p*), divided by the area of the rectangle, *AP*. **(b)** In all our extinction models the value of *R* depends on the order in which plants are made extinct, so many simulations are run with random sequences of primary extinctions to produce a distribution of robustness values *f*(*R*).

Extinction models have been adapted in various ways. One approach has been to use specific sequences of primary extinctions, such as ordered by traits of the nodes (Dunne, Williams & Martinez, 2002; Memmott, Waser and Price, 2004; Pocock, Evans & Memmott, 2012; and Santamaria *et al.,* 2016). A different adaption, developed by Vieira and Almeida-Neto (2015), allowed co-extinction due to feedback between guilds (implemented stochastically, based upon interaction frequencies), so permitting the possibility of cascades of extinctions. Traveset, Tur and Eguíluz (2017) created a variant of this model, incorporating empirically-estimated dependencies of plants on pollinators. Another development was to allow rewiring of edges (pollinators switching from one plant to another) based on empirical evidence (Kaiser-Bunbury *et al.* 2010). All of these aim, in some way, to increase the biological realism of extinction simulations - an issue that many of these papers acknowledge is lacking.

Early extinction models showed that the robustness of communities to random primary extinctions increased with network connectance, i.e. the fraction of the possible interactions that were actually observed (Dunne, Williams & Martinez, 2002) and the resulting robustness was often interpreted in terms of network nestedness (Memmott, Waser and Price, 2004). Vieira and Almeida-Neto (2015) found that cascades were more likely in highly connected networks. However, more detailed investigation of the impact of network structure on robustness has been lacking. These studies form the foundation of extinction models for plant-pollinator communities on which we base our work.

At first, many of the observed plant-pollinator networks were described with binary edge weights; interactions between pairs of species were either observed or not. Increasingly researchers are measuring the frequency or importance of interactions to create weighted networks, yielding a better description of the interactions observed (Ings *et al.,* 2009; Tylianakis *et al.*, 2010) and accounting better for under-sampling biases (Bersier, BanaŠek-Richter & Cattin, 2002). But the lack of weighted robustness models means that the information on the interaction weights, when known, either has to be discarded (e.g. Pocock *et al.* 2012), be used to weight the binary outcomes by node abundance (Kaiser-Bunbury *et al.* 2010) or be incorporated for plants only (Traveset, Tur & Eguíluz, 2017). Therefore, there is a need for robustness models that use all of the weighted data available. Here, we aim to address this need.

One of the features of these extinction models is that when using random sequences of primary extinctions on a single network, there is a broad distribution in the resulting robustness values (see Fig. 1b). Robustness is therefore a product both of structural heterogeneity of the network (eg Pastor *et al.*, 2012) and of the method of producing extinction sequences; we explore each of these contributions to robustness in this paper.

Here, our aim was firstly to develop a suite of extinction models which can be applied to weighted, as well as binary, bipartite interaction networks. Secondly, we paid particular attention to how robustness (assessed using our different models) is affected by network structure and how our models influence measured robustness. Throughout this paper we apply our models to mutualistic bipartite networks (specifically plant-pollinator networks) but discuss how they can be applied to any bipartite network (e.g. with trophic, uni-directional interactions).

## 2. MATERIALS AND METHODS

In this study, we examine the robustness of observed plant-pollinator networks that describe observed interactions between species in a community. A network has *P* plant nodes and *A* animal nodes, and contains *E* interactions between species, encoded in the *A* × *P* matrix *<u>M</u>*. Interactions may be binary (b) or weighted (w).

We illustrate our models and findings using a plant-pollinator network, with data collected by Memmott (1999), from Ashton Court, a site in Bristol, UK. We will refer to this as the Ashton Court (AC) network. This is a well-sampled network (Bluthgen, Menzel & Bluthgen 2006) with interactions recorded over a short period of time (1 month). The AC network is highly resolved: all plants were identified to species (P=25) and many pollinators were identified to species level (and the remaining pollinators identified to morphotype: A=79). *<u>M</u>*^*AC*^ contains 104 species, *E*=299, with connectance (proportion of realised interactions) of 0.151. Interactions in the AC network are weighted by the number of observations. The degree distribution of plant species is highly skewed as is often the case in plant-pollinator networks.

We also present results for two other networks to illustrate the generality of our model outcomes. These networks were collected in Shelfhanger (Sh), Norfolk, UK (Dicks, Corbet & Pywell, 2002); *<u>M</u>*^*Sh*^ has *P*=16, *A*=36, *E*=89 and connectance 0.148; and in Ottowa, Canada (Small, 1976); *<u>M</u>*^*Ot*^ has *P*=13, *A*=34, *E*=141 and connectance 0.319. Both networks have interactions weighted by the number of observations. We selected these networks because, like the AC network, they describe northern hemisphere, temperate ecosystems, and have a similar size to the AC network, but differ in having lower and higher connectance respectively.

### 2.1 Model Development

We took as our starting point the extinction model of Memmott, Waser and Price (2004), who analysed the robustness of binary networks by making species of one type (in their case, pollinators) extinct in a random order, i.e. they used a random primary extinction sequence. From this, we developed two new extinction models that each include sub-sequences of plant extinctions that are determined by network structure. All three models (summarized in Fig. 2) can be applied both to weighted and to binary interaction data by introducing a threshold rule.

**Figure 2:**
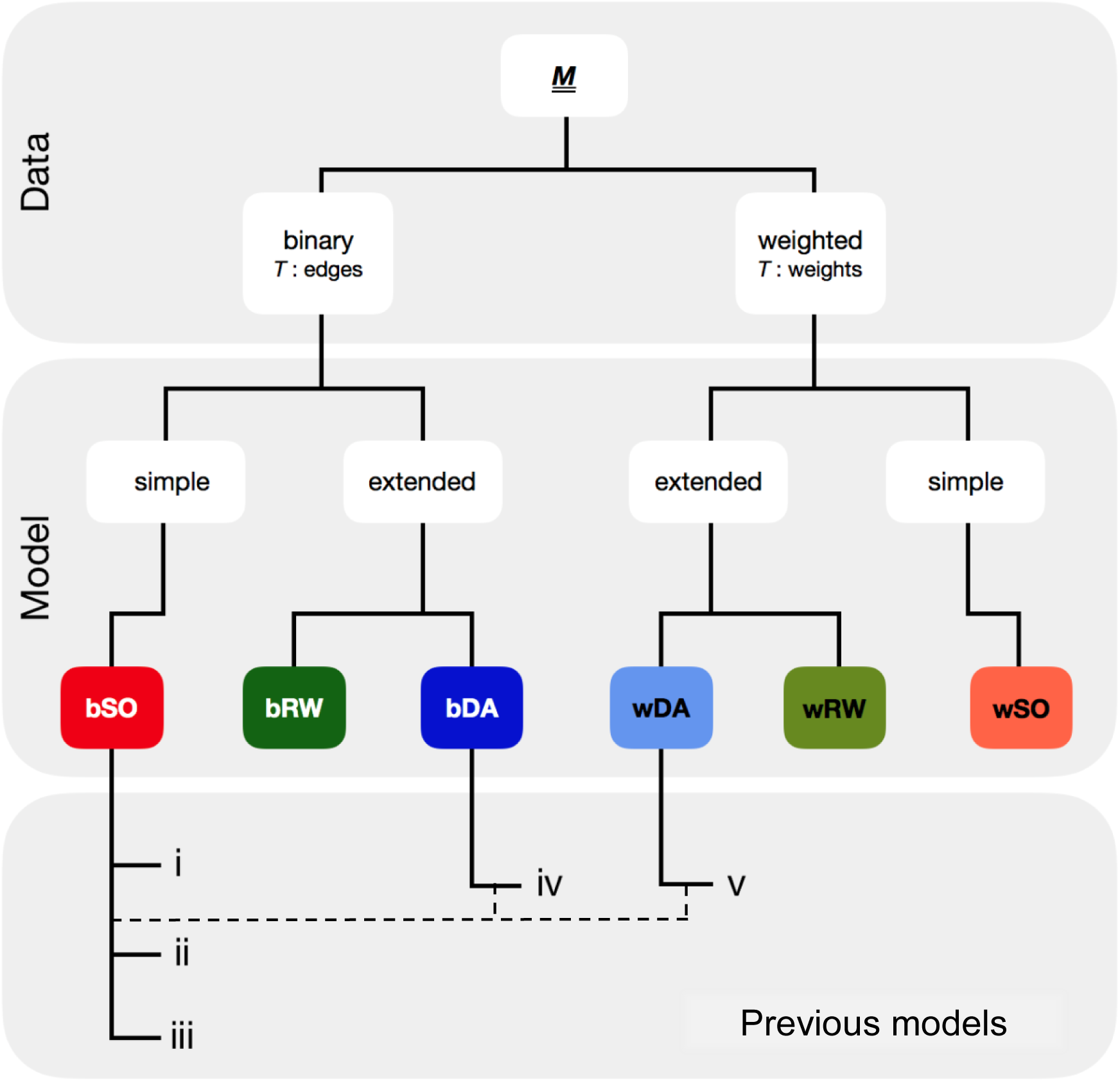
Framework of extinction models, with those used in this paper highlighted in colour. All models start from an observed mutualistic bipartite network *<u>M</u>* that can be binary (prefix b) or weighted (w). For binary data the threshold *T* is applied to the number of edges; for weighted data it is applied to the weights. The models are split into those that produce entirely random primary extinction sequences: Secondary Only (SO), and those that introduce other methods for determining extinction sequence: Deterministic Avalanche (DA) and Random Walk (RW). (i-v) indicate previous studies that represent special cases of the models in the framework where (i-iii) *T* = 1: i) Dunne, William and Martinez, 2002; ii) Memmott, Waser and Price, 2004; iii) Kaiser-Bunbury *et al.* 2010, iv) where *T* is applied stochastically and extinctions can ‘cascade’ (Vieira and Almeida-Neto, 2015) and. v) a hybrid of iv) and bSO with empirical plant dependencies (Traveset, Tur and Eguíluz, 2017).

In this section, we first describe the features that are common to all our extinction models and then outline the distinctive features of each, highlighting the relationships between ours and previous extinction models.

### 2.2 Universal Model Features

Starting from the observed matrix *<u>M</u>*, a node of one guild (plants) is removed as a primary extinction. Extinctions result in the loss of interactions from *<u>M</u>*, monitored in the ‘reduced’ matrix *<u>C</u>*. The loss of interactions may, according to the rules of the particular model, result in the secondary loss of nodes of the other guild (pollinators). In our new models (see below) the rules admit the possibility of each secondary pollinator extinction giving rise to further knock-on plant extinction(s). These plant extinctions cannot be considered ‘primary’, but will take their place in what we shall continue to refer to as a ‘primary extinction sequence’ of the *P* plant species.

All models proceed until all plant nodes are removed and all species - plants and pollinators - are extinct. The robustness (*R*) of the network is calculated as

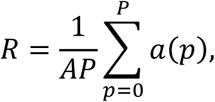

where *p* is the number of plant species that have gone extinct (from 0 to *P*) and *a*(*p*) is the number of pollinator species remaining in the network (from A to 0). *R* is the normalized (0 < *R* < 1) area under the curve of a graph of the proportion of plant nodes that have gone extinct against the proportion of surviving pollinator nodes (see Fig. 1a). Values of *R* closer to 1 indicate higher ‘robustness’ of the network to primary extinctions (Burgos *et al.,* 2007 and Albert, Jeong, & Barabási, 2000). We use *a*(*p*) as our response variable for all models in order to facilitate comparisons, although we note other options are possible: e.g. Kaiser-Bunbury *et al.* (2010) used the sum of interaction weights *w*(*p*).

The value of *R* is dependent on the specific sequence of primary extinctions, so running many random extinction sequences will, for all our models, produce a frequency distribution of values of *R* (Fig. 1b) which we denote *f*(*R*).

A key model feature we adopted is a threshold rule for secondary extinctions, so that a node becomes extinct once it has lost a fraction *T* or more of its observed interactions (binary *<u>M</u>*), or of its observed total interaction weight (weighted *<u>M</u>*). Clearly the value of *T* that we choose is arbitrary. It must lie in the range 0 < *T* ≤ 1. [*T*=0 is a pathological case; all pollinator species become extinct after the first primary plant extinction; *T* = 1 generates the extinction rule for most previous models (Dunne, Williams & Martinez, 2002; Memmott, Waser & Price, 2004; Kaiser-Bunbury *et al.,* 2010), although Vieira introduced a node-specific threshold ≤1.] We generated distributions of robustness *f*(*R*) for a range of threshold values (0.1 to 1 in 0.1 intervals) for three observed plant-pollinator networks (Ashton Court, Shelfanger and Ottawa) to determine the effect of *T* on the results of our extinction models. We then chose a threshold of *T* = 0.5 for the remainder of the paper: i.e. a secondary extinction occurs when a node has lost at least half of its interactions (binary *<u>M</u>*) or weights (weighted *<u>M</u>*). It should be noted that the ‘effective *T*’ (*T*_eff_) could be greater than *T*; for example, with a binary network and *T* = 0.5, a node linked to 5 others would go extinct after losing 3 edges, giving an effective *T* of 3/5=0.6. Since most pollinators are observed visiting a relatively small number of plants, the difference between the specified and the ‘effective’ threshold can be noticeable and so we calculate the node-averaged *T*_eff_ in all cases.

### 2.3 New extinction model features

We present three distinct models, which we denote: 1. Secondary Only (SO), 2. Deterministic Avalanche (DA) and 3. Random Walk (RW). Each model can be used with binary or weighted interaction data and is prefixed with ‘b’ or ‘w’ to indicate which.

#### Model 1. Secondary Only model (bSO and wSO)

In the Secondary Only model the order of primary plant extinctions is random. All pollinator extinctions are secondary and determined by the threshold rule. There is no spread of extinctions beyond the secondary extinction of pollinators. The method is as follows:

1. Select a random plant species (*e*) from those left in the network (matrix *<u>M</u>* the first time, then subsequently matrix *<u>C</u>*) for primary extinction
2. Make pollinator species connected to *e* extinct according to the threshold rule: if they have lost a proportion ≥ *T* of their original edges (bSO) or edge weights (wSO)
3. Count the number of pollinator species remaining, *a*(*p*), in the updated network (matrix *<u>C</u>*)

Repeat steps 1 to 3 until there are no species remaining in the network. Then calculate *R* according to equation 1.

In the special case *T*=1, the bSO and wSO models are identical to each other, and to the model described by Memmott, Waser and Price (2004). Kaiser-Bunbury *et al.* (2010) employed an adaptation to the special case *T* = 1 but used the weight of remaining edges *w*(*p*) as their response variable.

#### Model 2. Deterministic Avalanche Model (bDA and wDA)

In this model a randomly chosen primary (plant) extinction - a ‘trigger‘-may produce secondary extinctions (of pollinators) that themselves leave plant species with fewer than a fraction *T* of their observed interactions. If this happens, there is an ‘avalanche’ of plant extinctions. During the avalanche the sequence of plant extinctions is not random, but is determined by network structure. At the end of an avalanche a new, random, trigger is chosen. The method is as follows:

1. Select a random plant species (*e*) from those left in the network (*<u>M</u>* the first time, subsequently *<u>C</u>*) for primary extinction – this is a trigger 2.
2. 
  a. Make pollinator species connected to *e* extinct according to the threshold rule: if they have lost a proportion ≥ *T* of their original edges (bDA) or edge weights (wDA)
  b. Count the number of pollinator species remaining, *a*(*p*), in *<u>C</u>*
  c. Make plant species (there may be more than 1) extinct according to the threshold rule as above
  d. Repeat steps 2a to 2c until there is no further spread of extinctions, then repeat from step 1 with a new trigger

Repeat steps 1 and 2 until there are no species remaining in the network. Then calculate *R* according to equation 1.

Were *T* = 1 used here, step 2c would never result in tertiary plant extinctions and no avalanches would occur, so the DA and SO models would be identical. The ‘stochastic co-extinction model’ (SCM) developed by Vieira and Almeida-Neto (2015) is a special case of our bDA model where the threshold is applied stochastically and is node specific; specifically, extinctions of nodes at our step 2c occur with probability = 1-(remaining interactions) / (interactions at start). We adopt the term ‘avalanche’ for our spreading deterministic extinctions to differentiate them from the stochastic ‘cascades’ of Vieira and Almeida-Neto (2015), which occur once only, triggered by the first primary extinction. Traveset, Tur and Eguíluz (2017) employed what is essentially a hybrid SCM-bSO model, with empirical dependencies for plants and allowing only two-step cascades.

#### Model 3. Random Walk model (bRW and wRW)

The RW model is similar to DA, in that a trigger can cause an avalanche of non-random plant extinctions. This time, the order of plant extinctions within an avalanche is determined by the structure of the plant-plant projection network (described by the *P* × *P* matrix *F* which quantifies the number of pollinator species shared by each pair of plant species). The full method is as follows:

1. Select a random plant species (*e*) from those left in the network (*<u>M</u>* the first time, subsequently *<u>C</u>*) for primary extinction
2. Create the plant-plant projection network *F*
3. Select the next plant extinction (*f*) from the neighbours of *e* in *F* with a probability proportional to edge weights.
4. Make pollinator species connected to *e* extinct according to the threshold rule: if they have lost a proportion ≥ *T* of their original edges (bRW) or edge weights (wRW)
5. Count the number of pollinator species remaining, *a*(*p*), in the updated network (matrix *<u>C</u>*)
6. Identify plant *f* as the new *e* and make it extinct
7. Loop through steps 2 to 6. If no neighbours exist in step 3, revert to step 1.

Repeat steps 1 to 7 until there are no species remaining in the network. Then calculate *R* according to equation 1.

### 2.4 Natural extensions of our models

We have developed these three models for application to binary and weighted mutualistic bipartite networks and with a random order of primary plant extinctions (i.e. the selection of the next extinction in step 1 of Models 1,2 and 3 is random). However, we note that these models can easily be modified to use ordered primary extinctions, where the choice of plant in step 1 of Models 1,2 and 3 is according to a pre-determined rule (based on node degree, biological plant trait etc). The models can also be applied to bipartite networks with uni-directional dependencies (no feedback between the trophic levels, e.g. trophic or host-parasitoid interactions), though in that case avalanches cannot occur.

### 2.5 Comparison of robustness distributions from the three extinction models

The distribution *f*(*R*) generated from a single network *<u>M</u>* will depend on the model used and whether the data are weighted or binary. If there are *P* plant species in the network, there are *P*! distinct plant sequences. The SO models sample uniformly from these possibilities (i.e. all sequences are equally likely). The DA and RW models do not sample uniformly, because avalanches produce non-random sub-sequences determined by the structure of the network. Using binary and weighted versions of the Ashton Court (AC) network we generated 25,000 extinction sequences using each of the 3 models, in order to assess the effect of *R* on model choice. To create values of *R* that lie close to the theoretical maximum and minimum we ran bSO with plant extinctions in order of increasing and decreasing degree.

### 2.6 Assessing node and network-level determinants of variation in robustness

Having described the variation in robustness, we finally sought to assess the attributes which determine this variation under each model. We did this in two ways: by creating null versions of the AC network with defined characteristics, and by examining extinction rank (when a plant goes extinct in an extinction sequence). These calculations are designed to test the role of plant degree in determining the central tendency and spread of *f*(*R*) in our three extinction models.

#### Generating null networks to test effect of degree distribution

To explore the effect of degree distribution *g*(*k*) on robustness we created three exemplar null networks from the AC network (in binary form) with different degree distributions. Firstly, we calculated the expected (null) plant and pollinator degree distributions *g*_*E*_(*k*) from 10,000 random networks with the same number of plant species (*P* = 25), pollinator species (*A* = 79) and edges (*E* = 299) as the AC network. (Throughout we excluded networks with any disconnected nodes from further consideration.) We then generated 10,000 random networks from the AC matrix (*<u>M</u>*^*AC*^) according to 3 randomisation protocols; 1) randomise all values in *<u>M</u>*^*AC*^ to generate random plant and pollinator degree distributions; 2) randomise *<u>M</u>*^*AC*^ by row only so that the pollinator degree distribution is preserved; 3) randomise *<u>M</u>*^*AC*^ by column only so that the plant degree distribution is preserved. We selected, from each set of random networks, the exemplar network which had the best match to *g*_*E*_(*k*) for 1) both plants and pollinators, 2) only plants and 3) only pollinators. The best match was assessed using the sum of the absolute difference between *g*_*E*_(*k*) and the random network degree distribution for either plants, pollinators or both. The bSO, bDA and bRW models were run on the observed AC network and on these three exemplar random networks to produce 10,000 extinction sequences, and a corresponding distribution of robustness *f*(*R*) for each.

#### Examining plant extinction rank in relation to *R*

To explore whether (for example) high-degree plants tend to go extinct toward the beginning of a primary extinction sequence, we defined the position in a sequence when a plant became extinct as its extinction rank (*r*), 1 ≤ *r* ≤ *P*. We ran each extinction model 25000 times, using binary and weighted versions of the AC dataset, and compared *h*(*r*), the distribution of extinction rank for each species generated by the simulations. From *h*(*r*) for each species we calculated the median extinction rank (*r*_m_) and degree (*k*) and tested for correlation using the Spearman coefficient. We expected that, by definition, *r* would be equal across all species for the SO model but not for the DA or RW models, since avalanches and random walks will tend to select (or avoid) high-degree nodes preferentially.

### 2.7 Testing on other networks

We tested our models on the Shelfanger and Ottawa networks. For each network we generated 25,000 extinction sequences, using each of the 3 models, in binary and weighted form. We used a fixed threshold of *T* = 0.5 for all cases as we are not directly comparing the networks, only seeking to confirm the generalities of the resulting *f*(*R*) distributions.

## 3 RESULTS

### 3.1 Varying the value of the threshold for secondary extinctions

Median robustness *R*_m_ increases non-linearly with *T*, and the least robust of our three networks at low *T* becomes the most robust at high *T* (Fig. 3a). However, this appears to be an artefact of the relationship between *T* and *T*_eff_, the node-averaged effective threshold (Fig. 3b), because *R*_m_ increases linearly with *T*_eff_ and the three networks are increasingly robust in order of increased connectance, as found by Dunne, Williams and Martinez (2002), at all values of *T*_eff_ (Fig 3c). The remainder of our results are presented for the AC network only (where *T*_eff_ = 0.694 for our chosen *T* =0.5).

**Figure 3:**
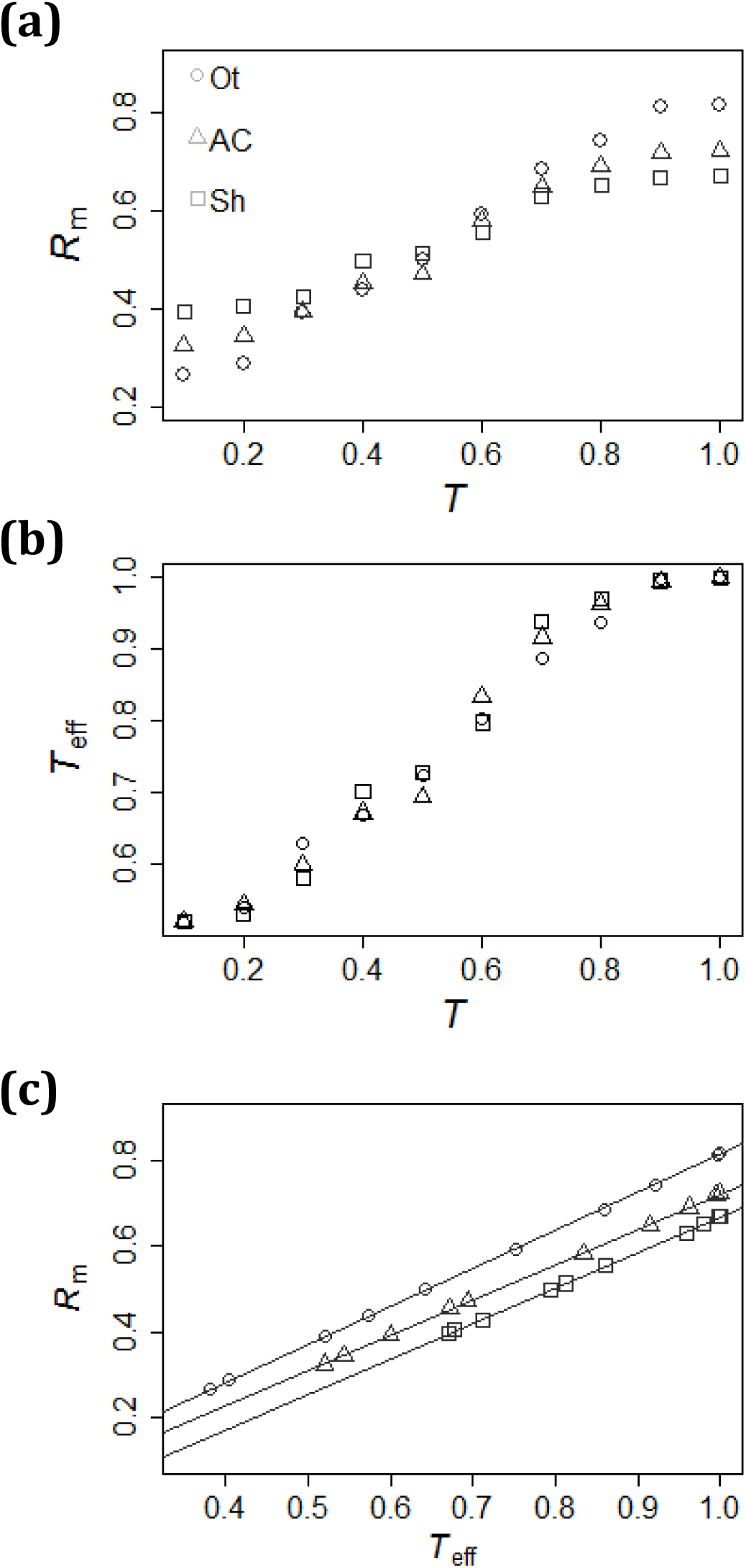
The relationship between selected extinction threshold (*T*), effective threshold (*T*_eff_) and median robustness (*R*_*<u>M</u>*_) for three plant-pollinator networks: Ashton Court (triangles), Shelfhanger (squares) and Ottawa (circles) using the bSO model. Variation of (a) *R*_*<u>M</u>*_ with *T*, (b) *T*_eff_ with *T*, and (c) *R*_*<u>M</u>*_ with *T*_eff_.

### 3.2 Robustness Distributions

The distributions *f*(*R*) produced by each of the 3 models for binary and weighted data (Fig. 4) are all rather broad, suggesting a strong dependence of *R* on the order in which plants are made extinct; the computed values span the range generated by primary extinction sequences in bSO with plants removed in increasing and decreasing order of degree (*R*=0.178 and *R*=0.812 respectively). The bSO model produces a relatively symmetrical *f*(*R*) with a median *R*_*<u>M</u>*_ *=* 0.473 and an inter-quartile range (IQR) 0.411 – 0.534. Using the bSO model as a baseline, the bDA model shifts *f*(*R*) to the right (Figure 4b: *R*_*<u>M</u>*_ *=* 0.512, IQR 0.439 – 0.588), inferring greater robustness, and bRW strongly shifts *f*(*R*) to the left (Figure 4c: *R*_*<u>M</u>*_ *=*0.369, IQR 0.330 – 0.411) inferring lower robustness. Using weighted, not binary, interaction data increases the IQR of *f*(*R*) in all cases.

**Figure 4:**
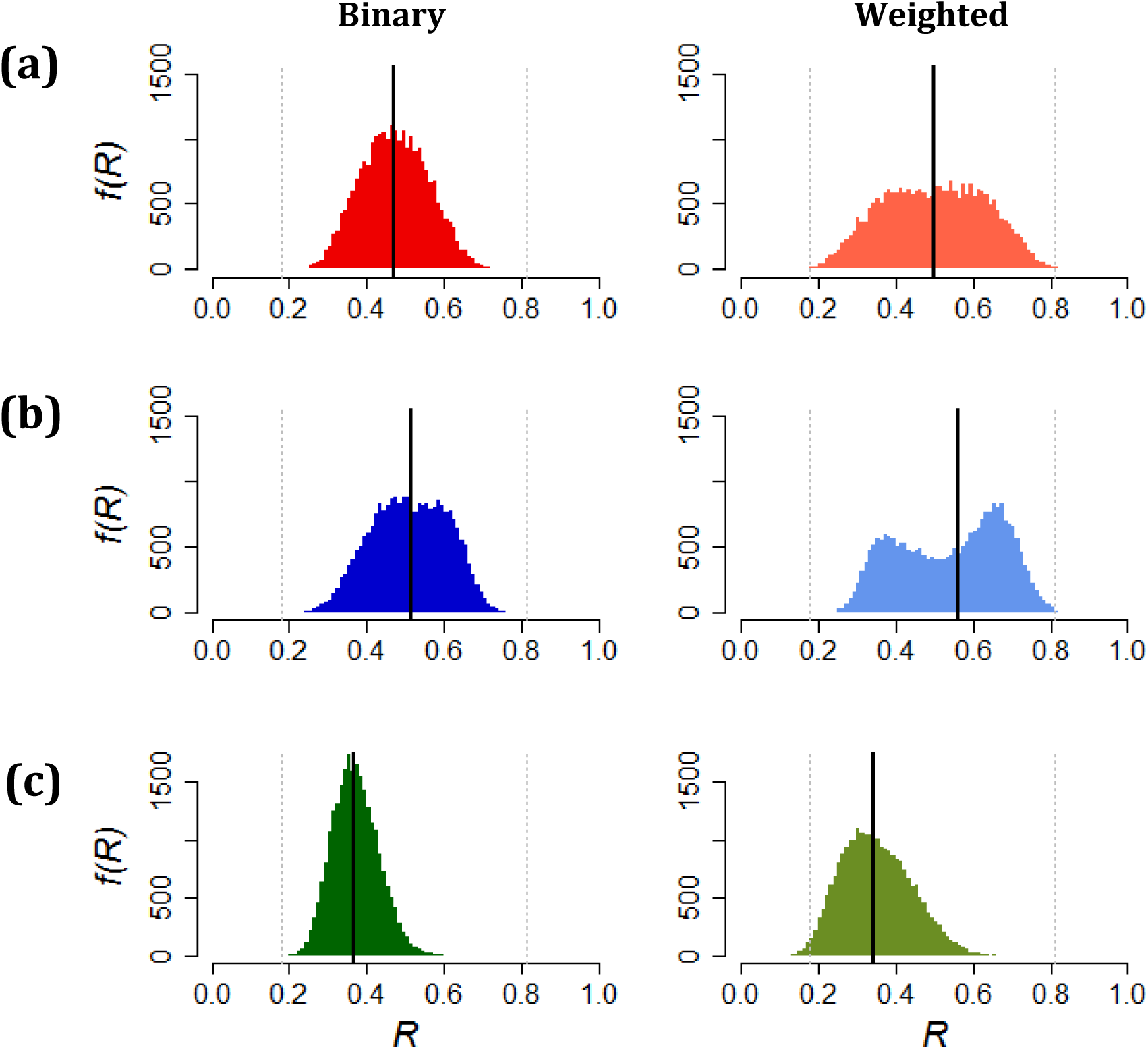
The distribution of robustness *f*(*R*) for the Ashton Court network, in binary (left column) and weighted (right column) form, generated by the three extinction models: (a) Secondary only (SO), (b) Deterministic Avalanche (DA) and (c) Random walk (RW). Median robustness *R*_*<u>M</u>*_ for each distribution is indicated by the solid vertical line. Dashed grey lines indicate *R* values for the bSO model generated by removing plant species in increasing (*R*=0.178) and decreasing (*R*=0.812) degree order.

### 3.3 Network randomisation test

Compared to the results of the binary extinction models for the AC network (Fig. 5a), we found that randomizing the degree distributions caused the distribution of robustness *f*(*R*) to be narrower (Fig. 5b-d), and this was especially so when the plant degree distribution is randomised (5c and 5d). This confirms that the observed, highly skewed, plant degree distribution of the AC network drives the broad robustness distributions we generate for this network. Note though that the *R*_m_ remain in the same order (RW<SO<DA) in every case, showing the consistency of effect from these models.

**Figure 5:**
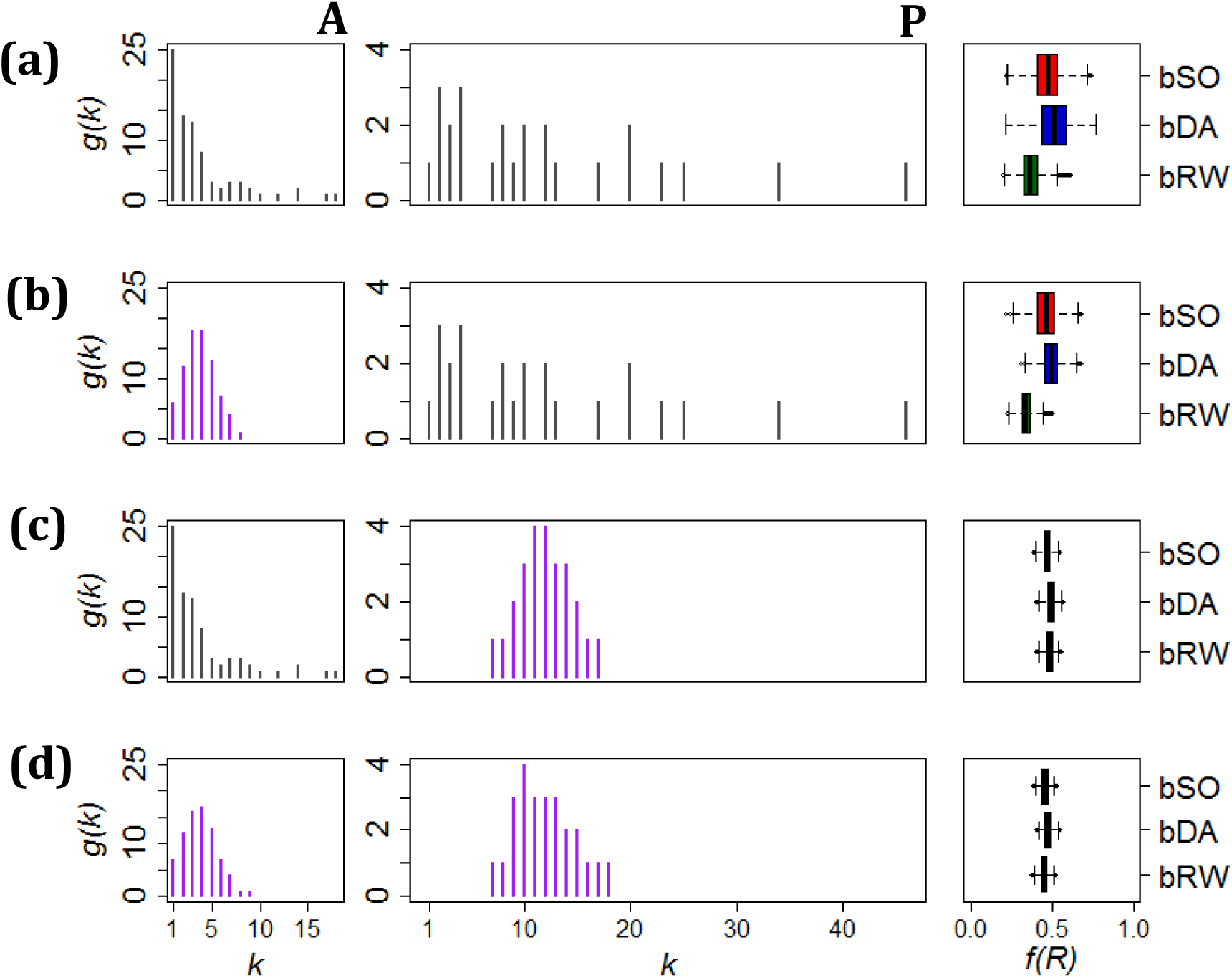
The effect of node degree distribution on robustness distribution *f*(*R*) for (a) the binary Ashton Court network and (b-d) randomised networks generated by the protocols described in section 2.6. Left column (A): pollinator degree distribution (grey - observed; purple – randomised); central column (P): plant degree distributions; right column: summaries of *f*(*R*) from the bSO, bDA and bRW extinction models. [Box-plots, with central lines showing median, boxes showing inter-quartile range, and whiskers showing the 95 % (2.5–97.5 %) interval].

### 3.4 Extinction rank of plant species, and the effect on *R*

Plant degree is a predictor of the plant’s extinction rank in the DA and RW models. In the SO models, the expected rank is constant for all plant species, irrespective of degree, because the extinction sequence is entirely random. In contrast, the observed extinction ranks of two example plant species from the DA and RW models are clearly skewed (shown in the insets of Fig. 6). In the DA models median extinction rank is positively correlated with plant degree (bDA: ρ = +0.803, P <0.0001; wDA: ρ= +0.464, P = 0.02). For the RW models, *r*_m_ is negatively correlated with *k* (bRW: ρ = –0.654, P = 0.0004; wRW: ρ = –0.723, P <0.0001). In other words, for the DA models, well-connected plants are resistant to extinction; the model preferentially prunes the low degree plants and network robustness is high compared to the SO models (Fig. 4b cf. Fig. 4a). In contrast, in the RW models plants with high degree are more vulnerable to extinction (the model preferentially ‘homes in’ on well-connected plants) which results in overall lower robustness of the network (Fig. 4c cf Fig. 4a). Clearly the apparent robustness of a network is dependent on the model used to incorporate feedback and structurally correlated co-extinctions.

**Figure 6:**
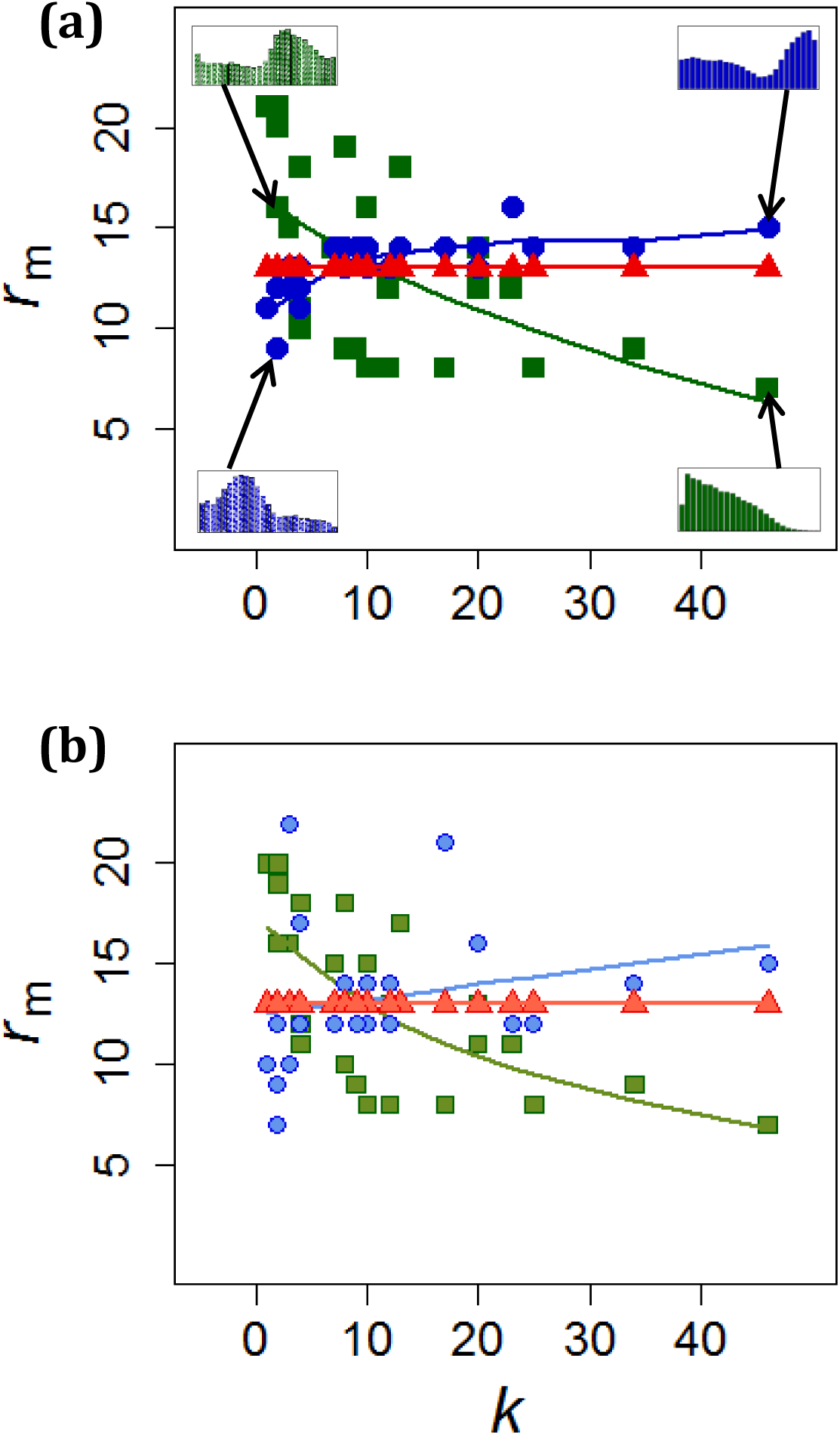
Variation of median extinction rank *r*_m_ with degree (*k*) for all 25 plant species in the Ashton court network for the three extinction models (SO: red, DA: blue and RW: green) and for (a) binary data and (b) weighted data. Smoothed lines are plotted as guides; Spearman’s rank correlation show that all these associations are significant: positive for DA (blue) and negative for RW (green). Insets illustrate the extinction rank distribution *h*(*r*) for two plant species (*Lathyrus pratensis* (*k*=2) on the left and *Daucus carota* (*k*=46) on the right) produced by the DA (blue) and RW (green) models. The corresponding median rank points are indicated with an arrow.

## 4. DISCUSSION

Robustness is a valuable quantitative metric used by network ecologists for describing and comparing the vulnerability of networks to simulated extinctions. We show, through the framework of extinction models that we developed here, that the measure of robustness is a consequence of both the model itself and the network structure.

Extinction models that calculate robustness have been around for a long time but until recently ecologists have used simple versions of these models (see Fig. 1). Here, we have created a framework of extinction models, building on those of Memmott, Waser and Price (2004), Kaiser-Bunbury *et al.* (2010) and Vieira & Almedia-Neto (2015), and applied it here to plant-pollinator networks. Importantly we have used an extinction threshold (pollinators can go extinct before all their plants go extinct and vice versa) that can be applied to all nodes. This addition has an ecological motivation - plants may decline to extinction due to declining pollination (as modelled by Traveset, Tur and Eguíluz, 2017) - and adds greatly to the flexibility of the model. Having *T*<1 allows us to create weighted versions of our models and provides the potential for feedback between the trophic levels and, hence, avalanches of extinctions cascading across the network (as first shown by Vieira and Almedia-Neto, 2015).

All of our extinction models, in binary and weighted form, produce a broad distribution of robustness values *f*(*R*) for each network that we analysed, indicating that there are aspects of the structure of the network that causes this variation. We found the degree distribution of the plants, in particular, to be an important driver of this variation. Plant pollinator networks tend to have fewer plant species than pollinator species (*P* < *A*), so the potential for a skewed plant degree distribution is increased, thus making it more influential on robustness in our test network (Memmott 1999).

Though ‘robustness’ has in the past been used to suggest priorities for conservation or management (Pocock, Memmott and Evans. 2012, Devoto *et al.* 2012), extinction models are not an attempt to predict precisely how an ecosystem would collapse. They do, nonetheless, offer a means to quantify and compare the structure of ecological networks, although to do this we need to ensure we are comparing like-for-like. One of the challenges is that robustness is influenced non-linearly by the value of the threshold *T.* However, we found that median robustness *R*_m_ is a linear function of the effective threshold *T*_eff_ and the effect is consistent across networks (Fig. 3). Therefore, if our models are to be used in the future to compare the robustness of ecological networks, it is important to note that *T*_eff_, not *T*, is used to ensure comparability.

Plant-pollinator communities are increasingly described with weighted interactions. We found (Fig. 4) that introducing weighted interactions has the effect of amplifying the outcomes observed for binary data: the inter-quartile range of the robustness distribution *f*(*R*) increases in all models for weighted networks, and the shifts in median robustness for DA and RW compared to SO are larger. These effects occur because weights tend to increase the skew of the plant degree distribution because high degree species accumulate high weights and low degree species only gain a small fraction of the overall weight in the network. The exaggeration of effects in *f*(*R*) highlights the importance of including interaction weights in robustness analysis, and in exploring all of the distribution *f*(*R*), not just its central tendency.

There are different ways in which feedback between trophic levels can be applied and we developed two illustrative models: the Deterministic Avalanche (DA) and the Random Walk (RW) models. These models (and others like the cascade model developed by Vieira and Almedia-Neto, 2005) may appear to be generating new outcomes but in reality, they simply produce a non-random sample of robustness values from those generated by a simple SO model. The Ashton Court dataset generated a huge range of *R* values, all of which can be realised in the Secondary Only models. The DA and RW models preferentially sample extinction sequences to produce skewed subsets of the SO outcomes (the *P*! extinction sequences are not all equally likely, and some will be impossible). The Deterministic Avalanche Model preferentially samples nodes that are 1 step away from each other in the network and extinctions can ‘fan out’ from each trigger (a randomly selected plant extinction). This mechanism can be likened to a ‘breadth first’ search on the network (see for example, Kolaczyk, 2009). The double-peaked distribution seen for the weighted DA model can be explained by competing effects: the model preferentially samples extinction sequences that have long avalanches and hence higher *R* values as well as those with shorter avalanches and hence lower *R* values, so the paucity of sequences with intermediate *R* values creates the two peaks. On the other hand, the Random Walk model follows a path though the plant projection network *F* away from trigger in the manner of a “depth first” search (Kolaczyk, 2009). Although both the DA and RW models are ecologically credible, they produce opposing results, demonstrating the influence of the model on the assessment of robustness. It is important for researchers using robustness models to have a clear justification for the model they use, and a clear understanding of how much their results are influenced by the model rather than the network data.

In ecological terms we can imagine the breadth first approach (DA models) as modelling the gradual collapse of the community along mutualistic dependencies. This is based on the assumption that the mutualistic relationship between plants and pollinators creates the potential for extinctions to travel in both directions between trophic levels (e.g. the presence and/or persistence of pollinators depends on flowers, and the presence and/or persistence of plants depends on pollinators). Avalanches in the DA model are only possible where dependencies are bi-directional, which can be perfect mutualism (each interaction is fully bi-directional) or it can represent a mixture of directed interactions (e.g. mutualism, nectar robbers, pollinator deception). The depth first approach (RW models) can be exemplified as a scenario where a plant disease is spread through the community by visiting pollinators, or a pollinator disease is spread through shared floral resources; a phenomenon observed by McMahon *et al.* (2015). The Secondary Only models are ecologically applicable when there is a uni-directional dependence in the interactions (e.g. presence of parasites depend upon hosts, but not vice versa) although, of course, they can be applied to mutualistic networks if dependence is assumed to be uni-directional (e.g. persistence of pollinators depends on flowers, but plants do not depend on pollinators).

All of these extinction models are designed to be applied to real ecological network data. Therefore, it is vital to consider the quality and reliability of the data being used. Empirical pollination networks vary hugely in sampling method, period of collection and taxonomic resolution, all of which can affect metrics of network structure. We caution against comparing the outcomes of extinction models across multiple networks, e.g. in meta-analyses or comparative analyses, without consideration of the data and the methods used to collect them. CaraDonna *et al.* (2017) highlight the potential pitfalls of assuming that a network constructed by aggregating samples over time is an appropriate representation of a community. Further work in understanding temporal variation and the description of fully-resolved plant-pollinator networks are key to improving the utility of extinction models.

Current robustness models lack the biological realism needed to make reliable ecological predictions. They are, however, useful for understanding and separating the effects of mechanism and network structure. We recommend therefore that researchers seeking greater ecological realism in models pay due attention to the details of the models themselves. Ecological conclusions drawn from robustness models may become less surprising when model developments are taken into account. We hope that by improving our understanding of extinction models at a mechanistic level, and by setting out different areas of model extension, our work will guide future developments in the analysis of the vulnerability of ecosystems to environmental change.

## ACKNOWLEDGEMENTS

We thank William Bonnell for support in developing the Random Walk model and the EPSRC for funding a studentship to MSB.

## AUTHORS’ CONTRIBUTIONS

All authors designed the methodology, discussed the results and commented on the manuscript at all stages. MSB coded the models and analysed the data with technical advice and support from RJ and MJOP. MSB drafted the manuscript. All authors contributed critically to the drafts and gave final approval for publication.

## DATA ACCESSIBILITY

This work has used the Web of Life database to access ecological network data: www.web-of-life.es. Model code will be deposited on the University of Bath Research Data Archive.

